# Cell-type specific burst firing interacts with theta and beta activity in prefrontal cortex during attention states

**DOI:** 10.1101/127811

**Authors:** B. Voloh, T. Womelsdorf

## Abstract

Population-level theta and beta band activity in anterior cingulate and prefrontal cortex (ACC/PFC) are prominent signatures of endogenously controlled, adaptive behaviors. But how these rhythmic activities are linked to cell-type specific activity has remained unclear. Here, we suggest such a cell-to-systems level linkage. We found that the rate of burst spiking events is enhanced particularly during attention states and that attention-specific burst spikes have a unique temporal relationship to local theta and beta band population level activities. For the 5-10Hz theta frequency range, bursts coincided with transient increases of local theta power relative to non-bursts, particularly for bursts of putative interneurons. For the 16-30Hz beta frequency, bursts of putative interneurons phase synchronized stronger than nonbursts, and were associated with larger beta power modulation. In contrast, burst of putative pyramidal cells were overall similarly beta-synchronized than nonbursts, but were linked with stronger beta power only when they occurred early in the beta cycle. These findings suggests that in the ACC/PFC during attention states, mechanisms underlying burst firing are intimately linked to narrow band population level activities, providing a cell-type specific window into the emergence, resetting, or termination of oscillatory activities.

## 1- Introduction

Narrow band population-level theta and beta band activity emerge during goal directed behavior in anterior cingulate and prefrontal cortex (ACC/PFC). The occurrence and strength of theta and beta activity thereby closely relates to behavioral functions including successful attention (Sheth et al. 2012; Voloh et al. 2015), correct retrieval of task rules (Buschman et al. 2012; Phillips et al. 2014), behavioral adjustment following errors (Womelsdorf et al. 2010), or short-term maintenance of stimulus-response mapping rules (Salazar et al. 2012; Babapoor-Farrokhran et al. 2017). These functional correlates of theta and beta activities emerge from the activation of cells and local circuits, but it has remained a fundamental open question which cell and circuit mechanisms are directly linked to these population level, narrow band activities (Kopell et al. 2014; Womelsdorf, Valiante, et al. 2014).

Growing evidence suggests that population level theta and beta activities are not supported equally by all neurons in a circuit, but rather that distinguishable cell-types show specific preferences to synchronize to the local electrical field at only a subset of narrow band frequencies (Hasenstaub et al. 2005; Roux and Buzsáki 2014; Womelsdorf, Valiante, et al. 2014). For example, in nonhuman primate prefrontal cortex, subsets of putative interneurons and putative pyramidal cells, defined by their narrow and broad action potential waveform shape respectively, show unique synchronization preferences to only beta or theta activity during attentive states in the primate (Ardid et al. 2015). Direct optogenetic control of spiking activity has likewise shown cell specific preferences to synchronize to the local oscillatory activity, with different subtypes of interneurons and pyramidal cells linked to theta, beta or higher frequency activity (Cardin et al. 2009; Sohal et al. 2009; Stark et al. 2013; Kim et al. 2015, 2016).

In addition to the cell-type, it has been documented that even for the same cell, not all spikes contribute similarly to population level rhythmic activity (Denker et al. 2011; Womelsdorf, Ardid, et al. 2014). This is particularly apparent for bursts, consisting of two or more spikes within a short (e.g. 5ms) time window. Previous studies suggests that bursts of pyramidal cells may directly index coordinated activity across larger recurrent networks (Larkum 2013). In particular, pyramidal cells bursts can be a direct consequence of coincident arrival of dendritic and somatic synaptic inputs from diverse distant sources (e.g. (Mainen and Sejnowski 1996; Larkum et al. 1999, 2007; Waters 2004; Manita et al. 2015; Sherman et al. 2016)). Bursts also have an outsized role in shaping neural activity; interneuron bursts can induce large compound inhibitory postsynaptic potentials strong enough to silence connected pyramidal cells (Hilscher et al. 2017), bursts of projection cells have enhanced postsynaptic efficacy in driving targets compared to singleton spike events (Swadlow and Gusev 2001), and induce more powerful longterm weight changes at their postsynaptic sites than singleton spikes (Birtoli and Ulrich 2004; Bittner et al. 2015; Wilmes et al. 2016). Moreover, it is believed that burst spikes are generated by mechanisms that are distinct from those of singleton spikes, suggesting that bursts could form a unique information channel during neuronal information processes (Krahe and Gabbiani 2004; Larkum 2013; van Ooyen and van Elburg 2014). Taken together, these characteristics assign bursts a particular role in the local neural circuit to shape how input is transformed into effective output of the circuit (Sahasranamam et al. 2016).

Despite the possible importance of bursts to shape network processes, direct support for the role of burst firing in coordinated network activity during actual cognitive processes is sparse. In a recent report we have documented that bursts firing of neurons in the ACC/PFC, but not isolated spikes of the same neurons, synchronized reliably to field activity in distant areas at narrow band theta, beta and gamma- band frequencies (Womelsdorf, Ardid, et al. 2014). This study pointed to burst firing events as a unique signature of long-range network activity, but left unanswered how bursts interact in the local circuits in which the burst event occurs.

Here, we first build on these earlier results and show that burst rate and the proportion of burst firing of neurons in ACC/PFC show sustained increases during a selective attentional state. These burst rate increases emerged in neural circuits whose population activity is characterized by power spectral peaks in the theta and beta band. To connect burst firing with population level theta and beta band activity, we characterized spike-triggered LFP activity around burst spikes and non-burst spikes. We found that spikes constituting the beginning of a burst firing event coincide with transient increases in theta LFP power when compared to non-burst spikes. This burst specific theta power modulation was particularly apparent for bursts of putative interneurons that were identified by their narrow action potential waveform. Independent of the theta power burst relationship, we found for the beta frequency band that burst spikes synchronized stronger to phases of the beta cycle than non-burst spikes, but without concomitant modulation of beta power. These findings reveal cell-type specific relationships of burst firing with meso-scale network activity indexed by narrow-band LFP components.

## Methods

### Experimental Procedures

Experiments were conducted in two awake and behaving macaque monkeys as described in detail in (Kaping et al. 2011), following the guidelines of the Canadian Council of Animal Care policy on the use of laboratory animals and of the University of Western Ontario Council on Animal Care. Extra-cellular field potential and action potential signals were recorded in each recording session from 1-6 tungsten electrodes (impedance 1.2-2.2 ΜΩ, FHC, Bowdoinham, ME) through standard recording chambers (19mm inner diameter) implanted over the left hemisphere in both monkeys. Electrodes were lowered through guide tubes with software controlled precision microdrives (NAN Instruments Ltd., Israel) on a daily basis, through a recording grid with 1 mm inter-hole spacing. Before the first recording session, anatomical 7T MRIs were obtained to visualize and reconstruct electrode as described in detail in (Kaping et al. 2011). Data amplification, filtering, and acquisition were done with a multi-channel processor (Map System, Plexon, Inc.), using headstages with unit gain.

The recording experiments were performed in a sound attenuating isolation chamber (Crist Instrument Co., Inc.) with monkeys sitting in a custom made primate chair viewing visual stimuli on a computer monitor (85 Hz refresh rate, distance of 58 cm). The monitor covered 36° × 27° of visual angle at a resolution of 28.5 pixel/deg. Eye positions were monitored using a video-based eye-tracking system (ISCAN, Woburn, US, sampling rate: 120 Hz). Eye fixation was controlled within a 1.4-2.0 degree radius window. Stimulus presentation, monitored eye positions and reward delivery were controlled via the open-source software MonkeyLogic. Liquid reward was delivered by a custom made, air-compression controlled, mechanical valve system with a noise level during valve openings of 17 dB within the isolation booth.

#### Behavioral task

Monkeys performed a covert selective attention, 2-forced choice discrimination task (**Fig. 1A**). Following a 2 second intertrial interval, a small gray fixation point was presented centrally on the monitor. Monkeys had to direct their gaze and keep fixation onto that fixation point until a change-event of the target stimulus late in the trial. After 300 ms fixation, two black/white grating stimuli were presented to the left and right of the center and contained oblique movements of the grating within their circular aperture. After 0.4 s each stimuli changed color to either black/red or black/green. After a variable time (0.05 to 0.75 s) the color of the central fixation point changed to either red or green, which cued the monkeys to covertly shift attention towards the stimuli that had the same color as the attention cue. Monkeys maintained central fixation and sustained covert peripheral attention on the cued stimulus until it underwent a transient clockwise or counter-clockwise rotation, ignoring possible rotations of the nonattended (uncued) stimulus, which occurred in 50% of the trials. In order to obtain liquid reward, the monkeys had to discriminate the rotation by making up- or downward saccades for clockwise/counter-clockwise rotations (the mapping was reversed between monkeys). Following this overt choice and a 0.4 s waiting period the animals received fluid reward (for a detailed description, see (Kaping et al. 2011)). A key component of the task is that the location of covert spatial attention on one of the two colored stimuli (left or right) is distinct from the possible locations to which the animal made a saccade (up or down) to indicate the transient rotation of the attended stimulus.

**Fig 1.**
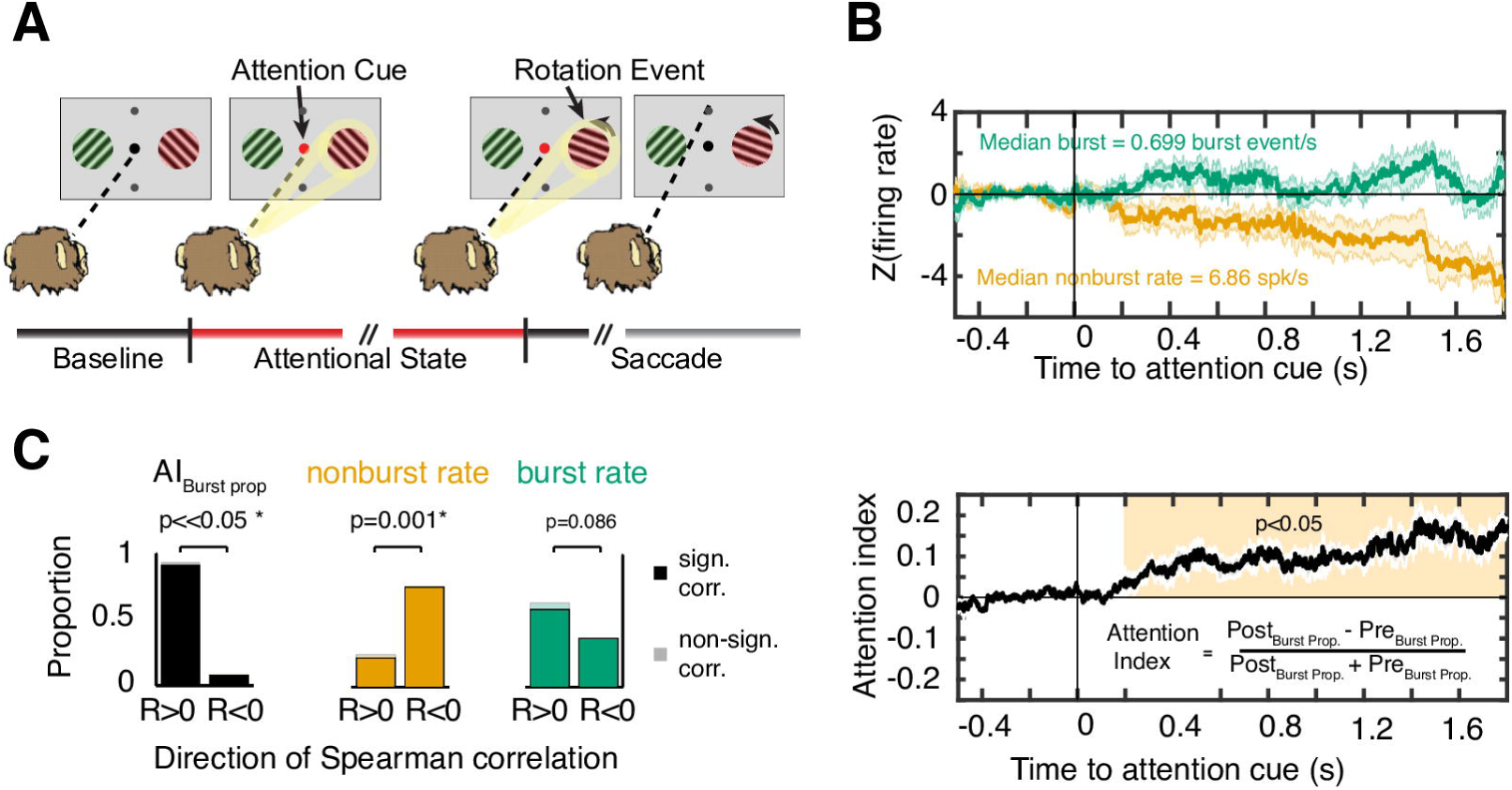
Attention task and increased proportion of burst firing following attention cue onset. (**A**) The task required continued central fixation starting in the Baseline Epoch (black). A color change of the fixation dot instructed to covertly shift attention to the color matching stimulus, marking the onset of an Attentional State (red bar). A rotation event in the cued stimuli had to be discriminated to receive reward by making an up-/downward saccade to clockwise/counterclockwise rotations (grey bar). Rotation events in the uncued stimulus (not shown) had to be ignored. (**B**) *Top panel*: Evolution of baseline-normalized burst rate (green) and non-burst rate (yellow) around the time of attention cue onset (n=41). Shading denotes standard error. *Bottom panel*: Burst proportion normalized relative to baseline, calculated as attention index. Grey shading denotes SE from bootstrap procedure and yellow shading shows time period with significant enhanced burst proportions (Wilcoxon signrank test, p<0.05). (C) Summary of monotonic trends in time. The proportion of cells that showed a monotonic increase (R>0) or decrease (R<0) when correlating the relevant variable with time. Transparent bars signify cells that did not reach significance individually. *Left:* 37/41 cells exhibit a monotonic increase in burst proportion after attention cue onset (χ^2^ test, p<<0.001). *Middle*: For 31/41 cells, the non-burst rate decreased in time (χ^2^ test, p=0.001). *Right*: Burst rate tended to increase in more cells after attention cue onset (n=24/41, χ^2^ test, p=0.08).

#### Neuron isolation

During recording, the spike threshold was adjusted such that there was a low proportion of multiunit activity visible against which we could separate single neuron action potentials in a 0.85 to 1.1 ms time window. Sorting and isolation of single unit activity was performed offline with Plexon Offline Sorter (Plexon Inc., Dallas, TX), using the separation of the first two to three principal components of the spike waveforms, and strictly limiting unit isolation to periods with temporal stability. For analysis we selected the subset of 422 maximally isolated single units whose waveform principle components were clearly separated with a density profile separated from the density profiles from multiunit background activity and other simultaneously recorded waveforms. The first two principle components explained on average 73.37% (± 1.3 SE) of variance across all waveforms that crossed thresholds. To quantify the separation of the waveforms’ first two principal component scores we calculated the Mahalanobis (ML) distance (using the Matlab function **mahal**). The ML distance metric uses the matrix of distances between datapoints to the mean, and the variance / covariance matrix to calculate the multivariate distances between points. We calculated the ML distance for the first two principal component scores of the spike waveforms of the recorded single unit relative to the scores of the waveform of the multi activity and noise of the same recorded channel and found an average ML distance of 24.12 ±1.8 (for examples see **Supplementary Fig. S1**).

#### Classifying cell types using spike waveform analysis

For all well isolated neurons we normalized and averaged all action potentials (APs) and extracted the peak-to-trough duration and the time of repolarization as described in detail in (Ardid et al. 2015). The time for repolarization was defined as the time at which the waveform amplitude decayed 25% from its peak value. Across the average waveforms of the cells we calculated the Principal Component Analysis and used the first component (explaining 84.5 % of the total variance), as it allowed for better discrimination between narrow and broad spiking neurons, compared to any of the two measures alone. We used the calibrated version of the Hartigan Dip Test (Hartigan and Hartigan 1985) to discarded unimodality for the first PCA component (p < 0.01) and for the peak to trough duration (p < 0.05) but not for the duration of 25% repolarization (p > 0.05). Additionally, we tested whether the distribution of the PCA score is better fit with two than one Gaussian. We applied Akaike's and Bayesian information criteria to test whether using extra parameters in the two-Gaussian model is justified. In both cases, the information criteria decreased (from -669.6 to -808.9 and from -661.7 to -788.9, respectively), confirming that the two-Gaussian model is better. We then used the two-Gaussian model and defined two cutoffs that divided cells into three groups. The first cutoff was defined as the point at which the likelihood to be a narrow spiking cell was 10 times larger than a broad spiking cell. Similarly, the second cutoff was defined as the point at which the likelihood to be a broad spiking cell was 10 times larger than a narrow spiking cell. This ensured across the whole population that 95% of cells (n = 401) were reliably classified: neurons at the left side of the first cutoff were reliably classified as narrow spiking neurons (18.7%, n = 79), neurons at the right side of the second cutoff were reliably classified as broad spiking neurons (76.5%, n = 323). The remaining neurons were left ‘unclassified’ as they fell in between the two cutoffs (4.7%, n = 20) (see **Fig. 2B**).

**Fig 2.**
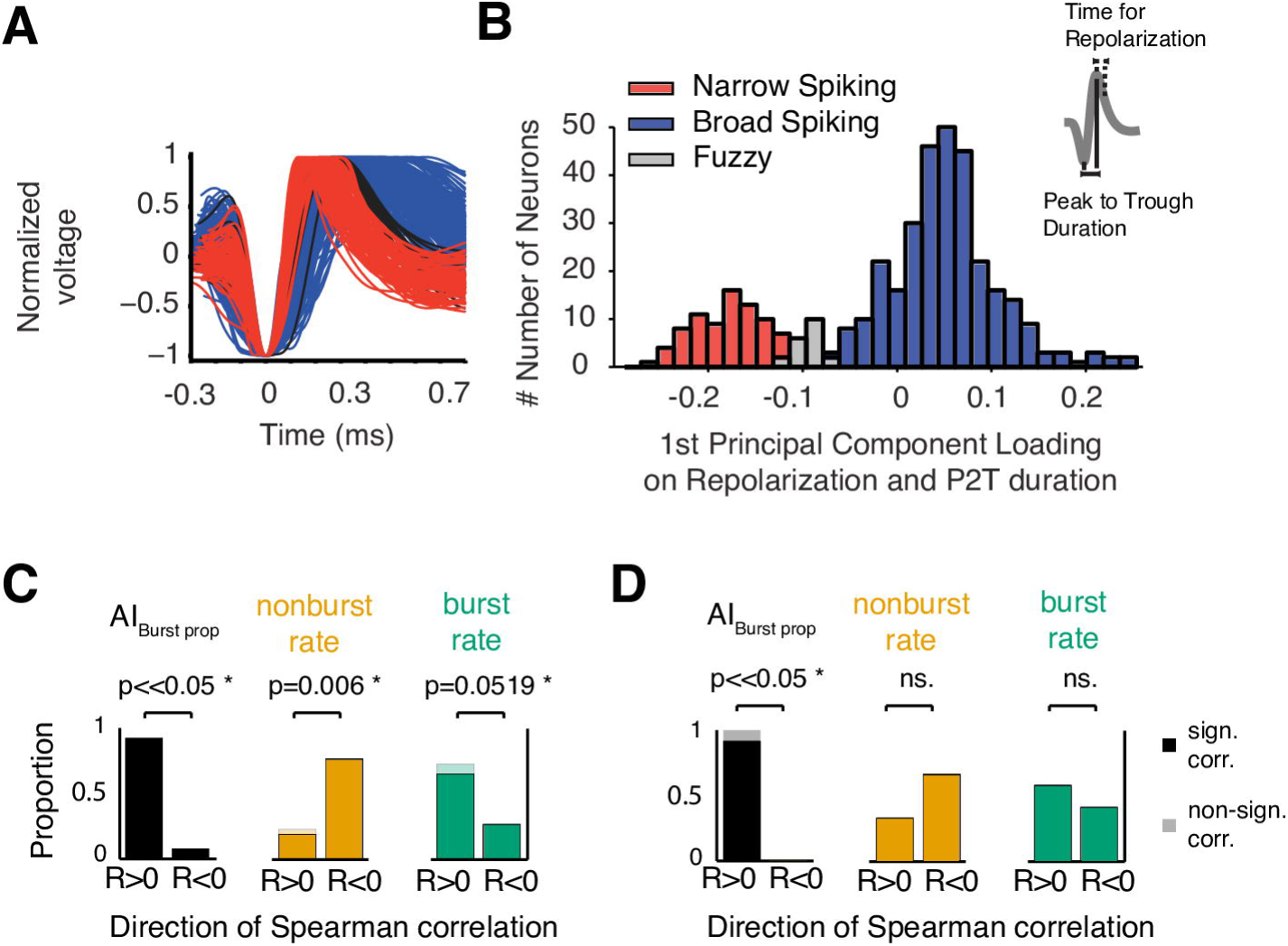
Cell-type specific modulation of burst and non-burst rate. (**A**) Average normalized action potential waveforms across all recorded narrow-spiking (NS, red), broad-spiking (BS, blue), and unclassified (grey) cells. (**B**) Bimodal distribution of NS and BS cells (and unreliably classified cells in between) as indexed by PCR score that combines the peak-to-trough duration and time to 25% repolarization of action potential waveforms (see inset). (**C,D**) Proportion of (C) BS and (D) NS cells with increased (R>0) and decreased (R<0) burst proportion (*left*), non-burst rate (*middle*), and burst rate (*right*) after attention cue onset relative to baseline.

### Data Analysis

Analysis was performed using matlab (The Mathworks). Throughout, we used conservative, non-parametric tests to draw our conclusions, and report throughout on the median. For the figures, the standard error of the median was estimated with a bootstrap procedure.

All analyses were performed on correct trials. The baseline period was defined as activity that occurred 0.5 sec. before cue onset. The attention period was defined as activity that occurred after cue onset, but before the first rotation of a stimulus, i.e. before target or distractor rotation (see **Fig. 1A**). To prevent experimental artifacts from affecting analyses, we ignored trials were the LFP deflection was greater than 10 SD away from the mean for that trial (median discarded trials = 0.78 +/− 0.002%). Bursts are rare events occurring less frequently than individual spikes. Moreover, low spike numbers can result in highly variable estimates of phase consistency (Vinck et al. 2010). For this reason, we selected only those cells that had a minimum of 30 burst events. To ensure sufficient number of spike/burst events for spectral analyses, we selected those selected neurons that had at least 30 burst spikes within the post-cue period with at least +/− 0.5 sec. of LFP data around the time of the spike. The +/− 0.5 sec. time window around spikes did never overlap with either the onset time of the centrally presented cue or the time of stimulus rotation. All LFP analyses were performed on the same channel as the spike.

To prevent spike artifacts in the LFP, we first lowpass filtered the LFP at 100 Hz using a two-pass 4th order Butterworth filter. Next, to prevent spike-locked artifacts, we used an interpolation approach when analyzing spike-triggered effects (Ardid et al. 2015). For each spike-centered LFP segment, a 5ms section was excised around the spike (thus including the second spike of a burst event), and we cubically interpolated over this segment.

To summarize, we selected cells that had (1) a minimum of 30 burst events and (2) had at least one second of LFP data around each event. This selection controls for any differences that might arise from changes in firing rate during attentional allocation (**Fig. 1**), and allows for an estimate of LFP power and phase synchronization using windows without possible transient onset responses to the cue or rotation events of the stimuli. That said, we recognize that this results in low cell numbers. To address this, we use non-parametric significance tests throughout so that the results are not biased by any outliers.

As an additional control to prevent artificial biases of spike LFP interaction, we consider spectral contents in the low frequency ranges (<30 Hz). This is to prevent, in particular, improper phase estimation that arises at higher (>30 HZ) frequency ranges (Ardid et al. 2015).

#### Burst definition

Burst events were defined as spikes that occurred with an interspike interval of ≤5 ms. All burst analyses were performed on the first spike of a burst event. To compute the burst proportion, the time-resolved firing rate of each cell was computed via the peri-stimulus time histogram (PSTH). The PSTH was computed separately for the baseline period (-0.5-0 sec.) and attention period from 0-2 sec. The PSTH was calculated with variable time windows (50, 100, 200, 300, 400, 700 ms). The choice of window length did not effect the main results. Time points with insufficient trials (n<=20) for an accurate estimate were discarded from analysis.

#### Time-dependent change in burst proportion

The burst proportion was defined as the portion of burst spikes relative to all spikes:

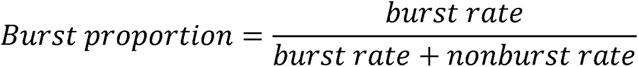

This procedure is intrinsically normalized for changes in the burst/non-burst rate that may occur (1) in time, or (2) across cells. The change in burst proportion during the post-cue attention state relative to the pre-cue baseline was calculated via a burst proportion Attention Index:

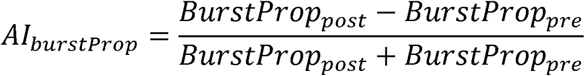

This was computed independently for each cell. The non-parametric Wilcoxon sign-rank test was used to determine if the burst proportion increases above baseline. The relationship between AI_burstProp_, non-burst rate, and burst rate with time was determined with the Spearman rank correlation, individually for each cell. These results were pooled to show if there was, on average, a monotonic increase (R>0) or decrease (R<0) in time. The overall trend (increase or decrease) was determined with a Chi squared test on the proportion of cells that showed either an increase or decrease.

#### LFP power analysis

We determined the dominant oscillatory components present in the LFP via spectral decomposition. For each trial, we set data after the time of the stimulus change to zero, thus analyzing activity only within the attention cue period void of possible on-responses to the rotation of the stimulus. We analyzed LFP power over a period [0.2, 2] sec after attention cue onset, thus preventing influence of transients related to directly after attention cue onset. Power was determined by Hanning tapering segments and performing a fast Fourier transform (FFT) over a range 2-40 Hz range. We scaled power by the frequency, in order to account for 1/f structure of LFP data, and normalized the range of each spectrum by [0 1] for comparison across cells.

To determine peaks in the spectral density plot, we used a peak detection algorithm based on Matlab’s findpeaks algorithm. First, we smoothed this plot to prevent the influence of noise. Next, we found peaks that were a minimum of 4 points away from each other, and that were above a threshold defined as 50% of the difference between the maximum and minimum of the individual spectra. This procedure extracted oscillatory components with a peak in the spectral density plot. We corroborated the output of this algorithm via a visual inspection of the spectral plots.

For further analysis of burst related power modulation and synchronization we defined the theta band as 5-10 Hz without including the 4 Hz bin in order to comply with our criteria that there should be at least 5 cycles of LFP data around each spike and 30 burst spikes minimum. Without this, more cells would not meet our criteria and thus reduce the pool of neurons for analysis.

#### Spike-triggered LFP power

We used the fieldtrip toolbox to get an estimate of the spectral content centered around each spike. For each LFP segment around the spike, we calculated power for frequencies ranging from 5-30 Hz, with 0.5 Hz steps, with an adaptive 5-cycle window per frequency. Signals were transformed with a Hanning taper before FFT. We then averaged the power of individual LFP segments locked to bursts or non-bursts individually. We then normalized the power across frequency and burst vs non-bursts to a range [0 1] (thus preserving relative difference in power across frequencies and spike types). We report on the median power across cells around bursts and non-bursts spikes. As well, we estimated the standard error of the median with a bootstrap procedure. We then determined if there was a difference in power depending on the spike type, with a non-parametric pairwise Wilcoxon sign rank test.

#### Analysis of time-resolved spike-triggered LFP power

To assess how bursts relate to oscillatory activity in time, we determined theta/beta power aligned to spike onset with a time resolved approach. We computed power as indicated above, but over a shorter 3-cycle adaptive window. This window was slid from -0.2 to 0.2 sec relative to spike onset with a 1 msec time step. Based on our previous results, we analyzed theta and beta effects separately, by averaging the power in a 5-10 Hz, or a 16-30 Hz band, respectively. To account for differences between cells, we Z-score normalized the time-resolved power (preserving differences in burst vs non-burst aligned power).

We assessed if there was a significant difference in time-resolved power relative to bursts or non-bursts with a Wilcoxon sign rank test. To correct for multiple statistical comparisons, we used a cluster based permutation approach) (Maris and Oostenveld 2007). First, we identified the largest significant cluster mass based on temporally adjacent stretches where p<0.05. Next, we shuffled the condition label and cell identity before recalculating the largest significant cluster. We performed this procedure 200 times, and compared the observed cluster against the permutation distribution. We then adjusted observed p-values of the points within a cluster according to the p-value of the cluster permutation test.

#### Phase synchronization analysis

To assess the degree of phase synchronization, we first extracted the angle of the individual LFP segments’ spectra for each frequency (from 5-30 Hz). Next, we identified cells that showed significant phase synchronized to a preferred phase by computing the Rayleigh statistic across all LFP segments (i.e. for both burst and non-burst spikes). Cells were considered to synchronize significantly at theta or beta if they had a significant frequency bin in the respective frequency range. We determined if the proportion of BS or NS cells that significantly synchronized was different with a Z-test for proportion, separately for the theta and beta bands.

To assess if bursts locked more strongly than non-bursts to particular phases, we used the Pairwise Phase Consistency (Vinck et al. 2010). This metric is not spuriously biased by differences in spike numbers. We computed the PPC for cells that synchronized significantly to either the beta or theta phases. To compare burst vs non-burst phase synchronization of the NS and BS cell populations, we averaged the phase consistency in the theta and beta bands, and assessed differences with the Wilcoxon signrank test.

The PPC can take on negative values for low sample numbers, which is uninterpretable (Vinck 2012). To this end, we converted raw PPC values to an effect size with the equation:

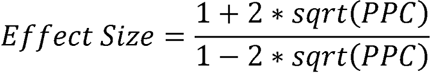

This effect size can be interpreted as the relative increase in spike rate at the cell’s preferred firing phase. For example, a PPC value of 0.01 corresponds to a 1.5 times greater spike rate at the preferred phase.

We determined the average theta and beta phases at which NS and BS spikes occurred by taking, for each cell, the average phase in the theta and beta bands. We report in the main text the mean and 95% circular confidence interval for NS/BS cells at theta/ beta, computed using the CircStats toolbox (Berens 2009). As well, to determine if NS and BS cells tended to fire at similar phases, we used the non-parametric Watson U2 two-sample differences in mean direction (Zar 2010), using the *watsons_u2_perm_test.m* function found online (http://www.mathworks.com/matlabcentral/fileexchange/43543-watson-s-u2-statistic-based-permutation-test-for-circular-data). Finally, to determine if bursts and nonbursts lock to the same phases, we used the circular median test on the pairwise difference in phases between bursts and nonbursts (Zar 2010). The null hypothesis was a median phase difference of zero, indicating the same preferred phase of firing for bursts and nonbursts across the population.

#### Phase-dependent power analysis

To identify the link between spike identity, phase of firing, and LFP power we calculated the phase-of-firing dependent power modulation. We began this analysis using the LFP spectra previously computed over a 5 cycle adaptive window (as described above). For each LFP segment, we then extracted the phase and power at each frequency. We next determined the preferred phase of firing of each cell (computed over all burst- and non-burst- aligned phases), subtracted this mean phase from the observed phases, and wrapped the transformed phases to the range [-pi pi], to obtain the phase relative to preferred phase of firing. This procedure allows comparison between cells, independent of individual cells’ preferred phase of firing.

Next, we binned the LFP power of each segment according to the phase at which they occurred (using 6 equally spaced phase bins). Results using different bin numbers (4, 5, 8, 9) were qualitatively similar. We then averaged the phase-binned power in the frequencies of interest, namely the theta and beta frequency band. This procedure was performed separately for segments around burst and non-burst spikes. Finally, we z-score normalized the phase-binned power across phase bins and spike-identity. As a first step, we determined if theta/beta was related to the phase at which the spike occurred. To this end, we took the median power in each phase bin across cell types, and fit a cosine of the form:

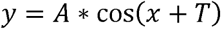
where *A* is the amplitude of the cosine, and *T* is the phase shift.

To determine whether power modulation was significantly different between bursts and nonbursts, we took the difference in power modulation between bursts and non-burst *(Ad = A_burst_ - A_non-burst_)*. We also assessed the difference in phase shift, first by converting phases into the time domain, and then taking the difference *(Td = T_burst_ − T_non-burst_)*. To assess significance, we randomly shuffled the spike identity and cell identity labels, recomputed the median power per phase bin, and refit the cosine. This procedure was repeated 1000 times to obtain a p- value for both the amplitude modulation, as well as the phase shift. This was done separately for BS and NS cells.

#### Testing for a relationship between spike-triggered LFP power and proportion of burst firing

To ascertain if the burst proportion was related to LFP power modulations, we correlated the burst proportion and theta/ beta power. The burst proportion was calculated for a 0-2 sec. period after cue onset. The average LFP power was calculated with a 5 cycle adaptive window (see above for more details) around the time of the (burst and non-burst) spikes. Power and burst proportion was z-transformed across cells. We used a Spearman rank correlation to determine the relationship between them. We ignored outliers, defined as power greater than 5 STD.

#### Relationship of spiketrain statistics and burst firing

To test whether intrinsic spiking properties of the cell are related to local theta and beta activity we quantified the burstiness of the spiketrain patterns of cells using the *local variability* measure (Shinomoto et al. 2009). Local variability describes the type of firing pattern that is most common to a neuron; LV values <1 indicate regular firing neuron, LV values ∼1 indicate a Poisson process; LV values >1 indicate an irregular/ burst firing. We correlated LV with average theta/ beta power with the Spearman rank correlation to determine if any power related effects that we see might be related to the cell’s propensity to burst.

## 2- Results

We recorded from different subfields of the rostral anterior cingulate cortex (areas 32 and 24) and lateral prefrontal cortex (areas 46, 9, posterior area 8), which we abbreviate as ACC/PFC, of two rhesus macaques performing a color-cued spatial attention task (**Fig. 1A**) (Kaping et al. 2011). Monkeys were cued to covertly shift attention to a target stimulus and sustain attention until the target stimulus transiently rotated clockwise or counterclockwise. Both monkeys performed the task well above chance (accuracy for monkey R: 75% STD: 8%; monkey M: 71%; STD: 11%) indicating that attention successfully shifted to the correct peripheral target stimulus following cue onset (Shen et al. 2014). We focus our analysis on correct trials and restrict analysis to the time immediately following attention cue onset and for the time in the trial before either of the peripheral stimuli transiently changed its motion direction. This period contained attention modulated neurons in all recorded brain areas as described before (Westendorff et al. 2016).

During task performance, we recorded 422 single neurons that were well isolated from background activities (**Supplementary Fig. S1**). To understand the relationship of burst firing (2 spikes within 5ms) to the local field potential during the attention state we extracted those 41 neurons (9.7%) firing at least thirty burst events across trials during the restricted time window of the selective attention state that was not influenced by strong onset responses to the cue or by transient changes of the peripheral stimuli (from 0.5 sec after cue onset and until 0.5 sec before the first motion change of a peripheral stimulus). 39% of these came from monkey M, and the remainder came from monkey R.

### 2.1 - Burst firing probability increases following attention cue onset

On average, bursts constituted 8.8% of all spike events. Across the 41 neurons, burst firing increased following attention cue onset, while the rate of non-burst firing decreased, rendering enhanced burst firing in ACC/PFC a signature of selective attention states (**Fig 1B,C**). Relative to the pre-cue baseline time period, the proportion of burst- over non-burst firing significantly increased from ∼200msec after attention cue onset (**Fig 1B**). 37 of 41 cells increased the proportion of burst firing during the selective attention state as indexed by Spearman rank correlations, which is a higher proportion than expected by chance (90%; χ^2^-test, p<<<0.05). The change in burst proportion was composed of both increases in the rate of burst firing (59%; **Fig 1C**; χ^2^-test, p=0.08), and decreases in non-burst firing (76%; **Fig 1C**; χ^2^-test, p=0.001). Neurons showing increased burst proportion following attention cue onset were similarly likely to show decreases, increases or no change in non-burst firing (χ^2^-test, p=0.48), suggesting that burst rate is modulated independently of non-burst firing rate.

Our extracellularly recorded neurons had action potential waveform shapes that reliably distinguished narrow spiking from broad spiking neurons based on their trough-to-peak ratio and their time to repolarization as reported previously (**Fig. 2 A,B**) (Ardid et al. 2015; Oemisch et al. 2015). Among the 41 cells selected for the burst analysis 12 (29.3%), 26 (63.4%), and 3 (7.32%) fell into the categories of narrow, broad, and unclassifiable neurons, respectively. This allowed analyzing burst rate changes separately for different neuron classes. Both, BS cells (**Fig. 2C**) and NS cells (**Fig. 2D**) significantly increased their burst proportion following attention cue onset. Increased burst proportions were composed of increases in burst firing, as well as decreases in non-burst firing rate (**Fig. 2C,D**).

### 2.2 - Relation of burst and non-burst spiking events to 5-10 Hz theta and 15-30 Hz beta band activity

We first asked how the attention-specific burst events relate to local field activity of the neural circuit. To answer this question, we first identified the frequency ranges in the LFP showing the most prominent oscillatory activity during the attention period of the task (0.2 - 2 sec.). We found that across the ACC/PFC, 74% (223/301) of the LFP recording sites had at least one clearly discernable power spectral peak in the theta or beta band indicative of periodically coordinated network activity (**Fig. 3A-D**). Of all recording sites, 25% (75/301) had power spectral densities with peaks within both, theta and beta frequency bands, 19% (58/301) of LFPs showed power peaks at the 5-10 Hz theta frequency range without concomitant beta modulation, and 30% (90/301) of LFPs showed only a power peak within the 15-30 Hz beta frequency band (**Fig. 3D**).

**Fig 3.**
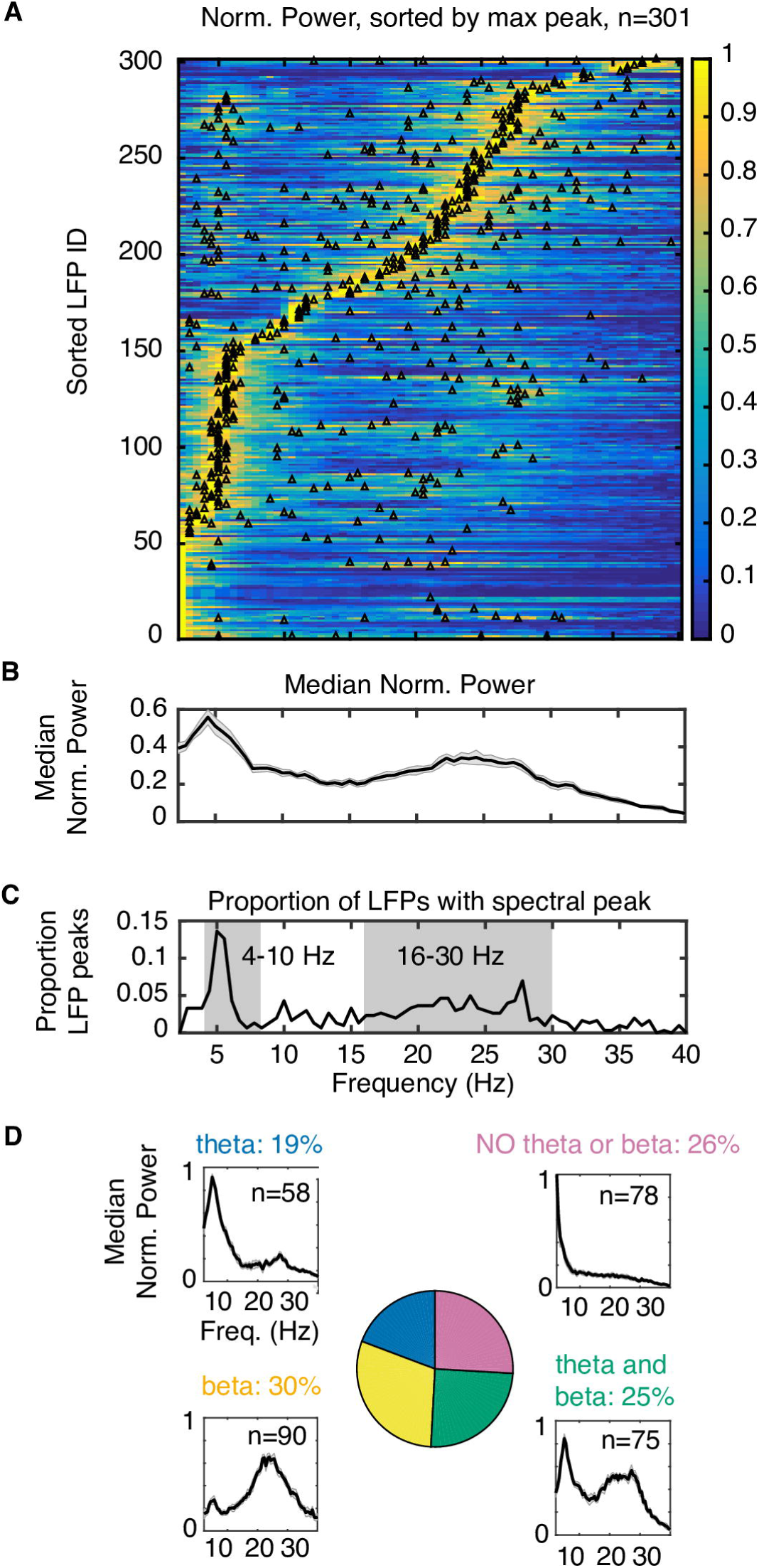
Theta and beta frequency components are the most prominent oscillatory signatures in LFP data. (**A**) Individual LFP power spectra (n=301, *y-axis*), sorted by the frequency of maximum power. Each power spectrum was normalized to account for 1/f noise, and scaled to the range [0 1] to compare across different LFPs. Triangles represent power spectral peaks with more than half-height amplitude (see Methods). (**B**) Median LFP power spectrum, revealing peaks in the theta and beta frequency range. (**C**) The proportion of LFPs that had a spectral peak at each frequency of interest. Grey shading highlights the theta and beta frequency bands. (**D**) Theta and beta peaks are both evident in 25% of recording sites, theta peaks without beta are evident in 19% of sites, and 30% of sites show only a beta peak without theta component.

The two main frequency ranges (theta and beta) with LFP power modulation were also apparent in spike-triggered LFPs across the 41 neurons recorded with sufficient number of bursts in the attention epoch (**Fig. 4A**; for an example of burst and non-burst spike triggered LFP see **Supplementary Fig. S2**). For the theta frequency band, the spike-triggered LFP triggered on the first spike of a burst showed significantly stronger theta power modulation than the spike-triggered LFP average for non-burst spikes (**Fig. 4B**; Wilcoxon signrank test, p<0.02). There were no power differences for bursts versus non-bursts in the beta frequency range.

**Figure 4.**
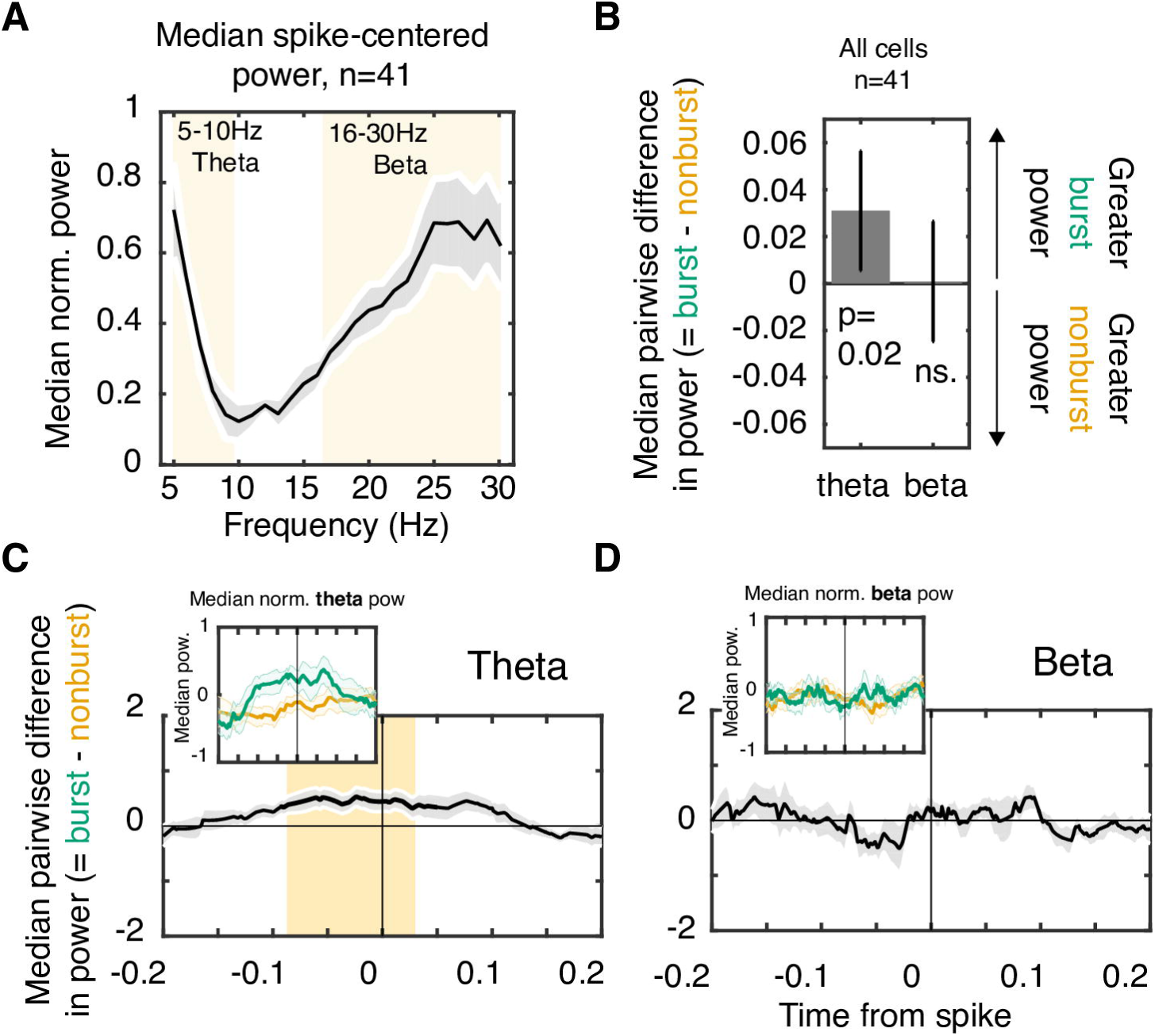
Theta and beta LFP oscillations are prevalent around spikes. (**A**) Median average LFP spectra around the time of spike occurrences. LFP power was controlled for *1/f^a^* noise and normalized to the range [0 1] before averaging. Shaded grey patches are the standard error determined with a bootstrap procedure, and the yellow background highlights the theta and beta ranges. (**B**) Median difference in spike-triggered average LFP power across all (n=41) cells in the theta and beta frequency band. Bars represent the standard error calculated with a bootstrap procedure. Theta LFP power is greater around bursts as compared to non-bursts, whereas beta power is equivalent. (**C**) Average pairwise difference in z-score normalized LFP theta power centered on bursts vs non-burst spikes. Significance at p <0.05 (orange shading) was assessed with a Wilcoxon sign-rank test, and multiple comparison corrected. Insets in the top left shows the time course of LFP power around bursts (green) and non-bursts (orange). Theta power is greater around bursts ∼26 ms before spike onset. (**D**) Same format as (*C*) for the beta frequency band, showing no average power difference between burst and non-burst spikes.

Spike-triggered LFP power modulation so far was calculated in symmetric ±0.5 sec. time windows around the time of the burst/non-burst event, leaving unanswered whether finer grained temporal analysis could reveal population level modulation preceding or following the burst/nonburst spiking events. To answer this question, we calculated LFP power in a sliding window analysis using adaptive 5 cycle windows every 1 msec around the time of the burst/non-burst event (see (Paz et al. 2008)). We found that across neurons, theta power was greater around burst spikes than non-burst spikes, starting as early as ∼80ms before the burst firing event (**Fig 4C**; p<0.05, Wilcoxon signrank test, multiple comparison corrected). There was no difference in beta power modulation for burst/non-burst spikes (**Fig 4D, Supplementary Fig. S3A**).

Enhanced theta power around burst events was visible in the subset of neurons with broad spiking waveforms (**Fig. 5A, Supplementary Fig. S3B**), but did not reach significance at a α=0.05 level. In contrast, burst events for the subset of NS cells were associated with significantly enhanced theta power when compared to non-burst spikes in a ∼110msec time window starting 26ms prior to burst onset (**Fig 5B**, p<0.05, Wilcoxon signrank test, multiple comparison corrected, see also **Supplementary Fig. S3C**). There was no consistent burst specific power modulation for BS or NS cells at the beta frequency band (**Fig. 5C,D**).

**Figure 5.**
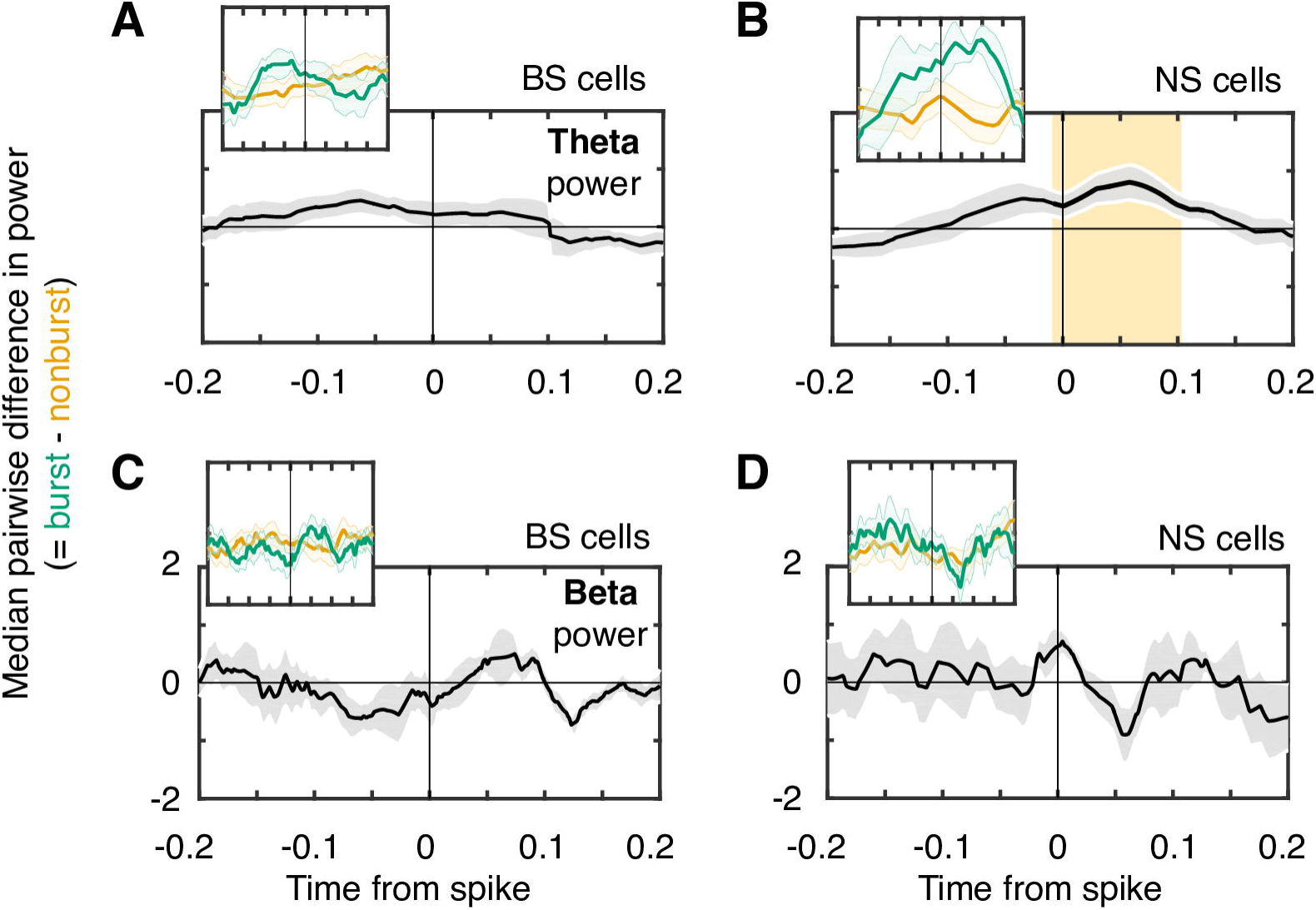
Time resolved LFP power around bursts of NS and BS cells. (**A,B**) Time resolved LFP power differences in the theta frequency and for burst versus non-burst spikes of (A) BS and (B) NS cells. Insets in the top left shows the time course of LFP power around bursts (green) and non-bursts (orange). BS cells (n=26) showed a nonsignificant increase in theta power before spike onset. NS cells (n=12) theta power significantly increased ∼57 ms after spike onset (shaded are). (**C,D**) Same as A,B for beta frequency power. There is no change in beta power around bursts relative to non-bursts. Error shadings denote SE computed with a bootstrap procedure.

### 2.3 - Burst specific spike-LFP synchronization in the beta frequency band

Spike-timing specific LFP power modulation indexes the strength of coherent network activity locked to the time of spikes, but does not indicate whether the spike events themselves synchronized to consistent phases of the narrow band LFP activity (Vinck et al. 2011). Across cells we observed multiple examples with significant phase synchronization at theta, beta, or both frequencies (**Fig. 6A**). Across the whole population, BS and NS neuron classes showed a large proportion of neurons with significantly phase synchronized burst- and non-burst spiking at the theta band (10 (83.3%) NS cells, and 14 (53.3%) BS cells. The proportion of theta locked neurons were not different at an alpha 0.05 level Z-test, p=0.08), as well as the beta frequency band (10 (83.3%) NS cells, and 16 (61.5%) of BS cells. The differences in proportions of NS and BS were not statistically significant, Z-test, p=0.18). NS and BS cells that showed spike-phase synchronization tended to synchronize to similar phases in the theta band (43.7± 90^0^ CI, 79.4 ± 90^0^ CI, respectively; Watson’s U2 test, p=0.9) and the beta band (-138 ± 39.8^0^ CI, -144 ± 48.7^0^ CI, respectively; Watson’s U2 test, p=0.279).

**Figure 6.**
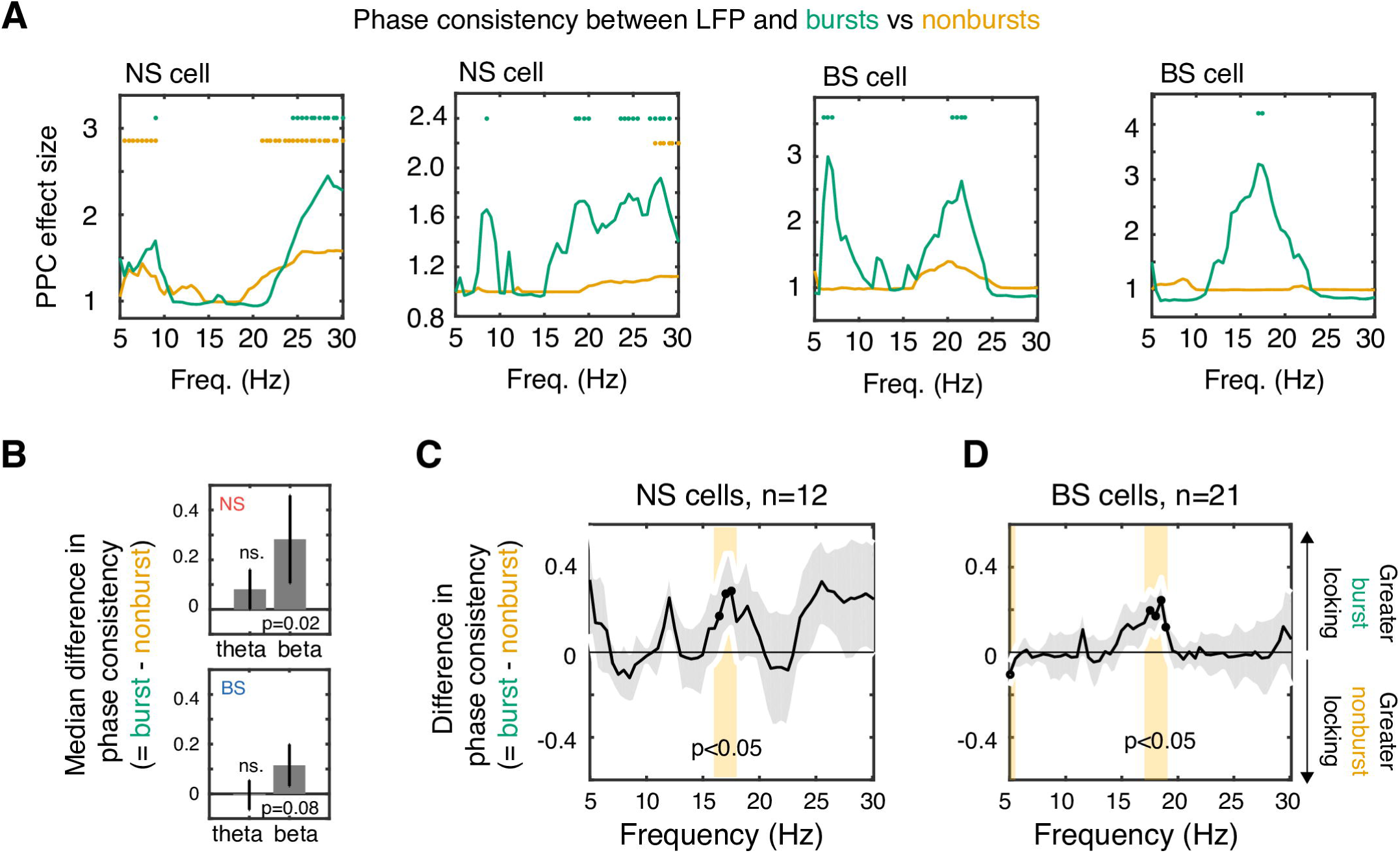
Increased phase locking to bursts is specific to the beta frequency band. (**A**) Examples of bursts (green) and nonbursts (orange) phase locking to the local activity. Significant locking (Rayleigh test at alpha<0.05) are marked with the appropriately colored dots. Note that even when nonbursts lock to theta or beta activity, locking to bursts is nonetheless stronger (compare panel 1 from 2). Beta-locked bursts tended to occur in a ∼15-20 Hz range, though was also apparent in a higher >20 Hz range in NS cells (panel 1, 2). (**B**) Median difference in phase locking in the theta and beta band, for NS cells (*top panel*) and BS cells (*bottom panel*). Burst events of NS cells synchronize stronger to the beta frequency than nonburst spike events. A similar trend exists for BS cells. (**C,D**) Median differences of burst versus non-burst to LFP phase locking for (C) NS cells and (D) BS cells. NS and BS cells show a narrow beta frequency band with significantly enhanced burst LFP locking over non-burst LFP locking.

For cells that did show significant phase synchronization at beta frequencies, burst spikes showed stronger synchronization than non-burst spikes (**Fig. 6B**). Across the broader 16-30 Hz beta band bursts of NS cells showed significantly enhanced burst-LFP synchronization compared to non-burst-LFP synchronization (n=11, p = 0.02, Wilcoxon signed rank test). A similar statistical trend was visible for BS cells (n=16, p = 0.08, Wilcoxon signed rank test; **Fig. 6B**).

Finer grained spectral analysis revealed for both, NS and BS cells, significantly enhanced burst over non-burst phase synchronization in a narrow ∼16-18 Hz frequency range (**Fig. 6C,D**). Moreover, analysis of the pairwise difference in phases of bursts and nonbursts showed both tended to synchronize at similar beta phases, both in NS cells (-135^0^±39.6 CI, -172^0^±47.7 CI for nonbursts and bursts, respectively; Median test, p=1) and (to a lesser degree) in BS cells (-130^0^±52.3 CI, -175^0^±50.4 CI for nonbursts and bursts, respectively; Median test, p=0.08) (*see also*, **Fig 7G**).

**Figure 7.**
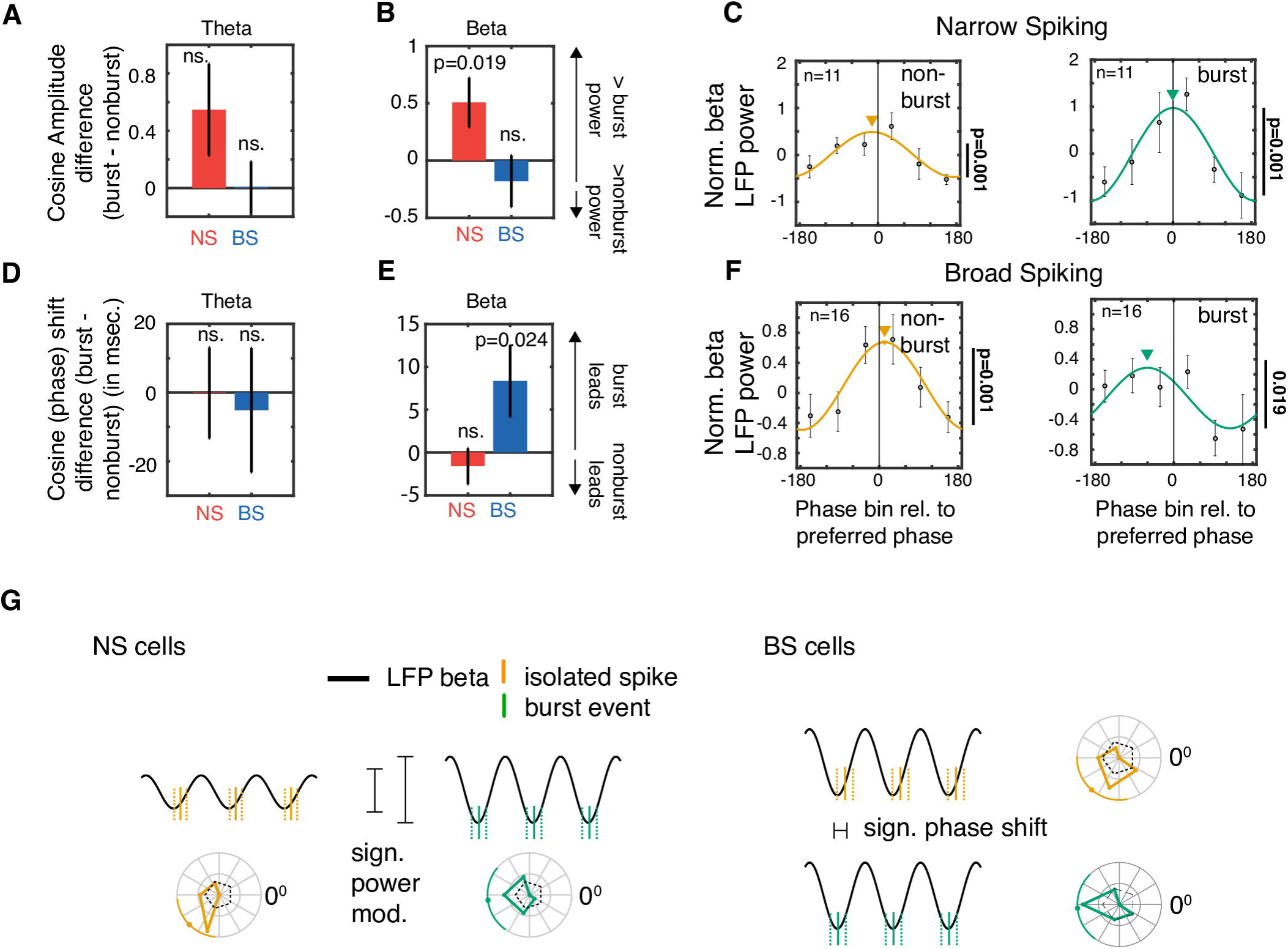
Phase-dependent power modulation of burst and non-burst events. LFP power as a function of the phase of firing of burst and nonbursts, for only those NS and BS cells that showed significant phase locking in the respective frequency band. (**A**) Theta LFP power is similarly modulated by the phase for burst and non-burst spikes for both NS and BS cells. (**B**) Beta power is stronger modulated by the bursts than non-burst LFP phases for NS cells, but not for BS cells. (**C**) Power (y-axis) significantly varied with the phase (*x-axis*) of non-burst spikes (*left panel*) and burst spikes (*right panel*) of NS neurons (randomization test, p<0.05), relative to the preferred phase of firing. Peak power for spikes synchronizing near their preferred LFP phase. (**D**) Difference (burst vs. non-burst spike LFP phases) of the phase (in milliseconds) at which power is maximal within a theta cycle, relative to the preferred phase of firing. The phase of maximal power modulation coincides with the preferred theta phase for both, NS and BS cells. (**E**) For the beta frequency band, bursts of BS cells that occurred prior to the cells preferred phase are associated with the maximum LFP beta power. The phase difference corresponds to ∼5-10 msec. (**F**) Same as (**C**), but for BS neurons. Beta power is significantly modulated by the phase (randomization test, p<0.05). Additionally, the burst-LFP phase with maximal power is significantly shifted relative to the preferred phase, preceding the preferred phase by ∼57 degrees (randomization test, p<0.05). (**G**) Summary sketch of beta modulation by burst specific phase of firing. Rose plots indicate the distribution of preferred firing phases – split by cell type and bursts vs singleton spikes – as well as the mean phase and 95% circular confidence intervals. *(Left panel)* In NS cells, both bursts and nonbursts occur near the same phase, but only the former leads to an increase in local beta power. *(Right panel)* On the other hand, in BS cells, for the same amount of beta power, bursts occur earlier in the cycle compared to nonbursts.

### 2.4 - Beta-band burst synchronization shows significant phase-dependent power modulation

The previous analysis showed that bursts synchronized stronger than non-bursts to beta-rhythmic but not to theta-rhythmic LFP fluctuations, while overall spike-triggered LFP power was stronger for bursts in the theta band, but not in the beta band. This dissociation of phase synchrony and power modulation could entail that LFP power around the time of the burst varied independently of the phase at which the burst and non-burst spike occurred in the theta and beta cycles (Canolty et al. 2012). To test this possibility, we calculated the *phase-dependent power modulation* by grouping burst spikes and non-burst spikes into six non-overlapping LFP phase bins. For each cell with significant phase synchronization, we defined its average spike-LFP phase as the preferred phase of firing, and averaged LFP power for each of the six phase bins across cells (Womelsdorf et al. 2012). This analysis showed that in NS cells, burst spikes were significantly stronger modulated by the phase in the beta frequency band than isolated non-burst spikes (**Fig. 7B,C**; randomization test, p=0.019). In contrast to NS cells, bursts of BS cells show similar strength of phase dependent power modulation than non-burst spikes (randomization test, p=0.43) (**Fig. 7B,F**). However, maximal beta power around bursts on average significantly led that of nonbursts (randomization test, p=0.024; **Fig. 7E**). This was evident as a shift of the phase (relative to burst onset) that contained maximal LFP power to 5-10 msec (57.5^0^) prior to the cells preferred phase of firing (**Fig. 7F**). In contrast to these burst specific effects in the beta band, phase dependent power modulation in the theta band was similar for bursts and non-burst and NS/BS cell classes (**Fig 7A,D; Supplementary Fig. S4**). In summary, these results show that at the beta frequency band burst firing is associated with prominent local LFP power similarly to the effects in theta band, but the burst effect at beta depended on the phase at which the burst and non-bursts occur within the beta cycle (*for a graphical summary:* **Fig 7G**).

### 2.5 - No correlation of power modulation and overall burst firing

Results so far characterized those bursts occurring during the selective attentional state following cue onset (0.5 sec. after cue onset and 0.5 sec. prior to a change of either target or distractor stimulus). It might thus be possible that burst mediated LFP power modulation, or the burst phase locking is particular prominent in neurons that show relatively larger modulation of burst firing. However, correlations of LFP power at theta or beta band did not correlate with burst proportion (**Supplementary Fig. S5**).

### 2.6 - Burst-specific modulation is not apparently linked to intrinsic neuron properties

The burst specific association with theta power and beta phases could be the result of neuron specific properties, or they may be better understood as network phenomena that emerge during active states irrespective of the intrinsic propensity of neurons to fire bursts. To address this issue, we computed the relative burstiness of the neurons spiketrains using the *local variability (LV)* metric that yields higher and lower values the more bursty (*LV*>*1*) or more regular (*LV*<*1*) the firing pattern is (Shinomoto et al. 2009). We found that NS cells (n=12) exhibited an average LV of 0.751 +/− 0.07 SE, and BS cells (n=26) showed 1.09 +/− 0.146 SE, which reflect average values for the overall recorded cell population (Ardid et al. 2015), suggesting that the selected BS and NS neurons are not intrinsically bursty neurons. Similarly, intrinsic firing rate variability was not a predictor of the overall degree of local oscillatory power in the theta band (Spearman correlation, R=-0.08, p=0.67) or beta band (R=0.10, p=0.529).

## 3 - Discussion

Here we reported that neurons in ACC/PFC increased burst firing proportionally to non-burst firing when nonhuman primates engage in a selective attentional state. Burst firing events were a signature of covert attention for both, narrow and broad spiking neurons. During the same attention state, we found that three quarters of recording sites showed prominent local field potential oscillatory peaks at a 4-10 Hz theta and/or a 16-30 Hz beta. Burst firing during the attention state had unique relationships to both theta- and beta-band population level activities. Within the theta frequency band, burst firing coincided with stronger theta power than non-burst firing. This theta effect was evident for bursts of broad spiking neurons, but it was strongest for bursts of narrow spiking neurons. Within the beta frequency band, bursts of narrow spiking neurons were stronger synchronized to the phases of the beta cycle than isolated spikes of the same neurons. For broad spiking neurons, burst spikes were associated with strong beta power at phases preceding the preferred phase. This result contrasted to non-burst spikes that showed a cosine shaped drop off in power away from the preferred phase (**Fig. 7F**). In summary, these results identify bursts as major signature of attentional states in nonhuman primate ACC/PFC and reveal burst specific modulation of local circuit field activity at those two oscillatory frequency bands that are closely associated with goal-directed controlled behavior (Phillips et al. 2014; Voloh et al. 2015; Babapoor-Farrokhran et al. 2017).

### 3.1 - Increased burst firing in ACC/PFC characterizes attention states and long-range activated networks

We found that burst firing in ACC/PFC increased shortly after a cue instructed subjects to deploy covert selective attention. The rate of burst firing rate increased at the same time as the firing of non-burst spikes decreased. This pattern of results renders ACC/PFC burst firing a unique characteristic of selective attentional processing states. This finding is consistent with the belief that dendritic mediated burst firing is a reflection of neurons within circuits participating in recurrent network activity in larger brain networks (Larkum 2013). According to this hypothesis, burst firing follows from coincident feedforward and feedback type synaptic inputs impinging on peri-somatic and distal dendritic regions of the burst firing neurons (Siegel et al. 2000; Larkum et al. 2004; Larkum 2013). Recent in-vivo experiments have begun to support this hypothesis of coincident distal and dendritic activation to underlie a unique processing state reflected in burst firing patterns (Manita et al. 2015; Palmer et al. 2016). During attentive states characterized by large-scale network coordination, enhanced dendritic activation in ACC/PFC could be the consequence of distant cortico-cortical axonal inputs, while perisomatic input could reflect prominent synaptic input from the thalamus and other subcortical sources (Barbas and Zikopoulos 2007; Miller and Buschman 2013; Barbas 2015; Womelsdorf and Everling 2015). We believe that our findings add critical support for the hypothesis that enhanced burst firing indexes an active recurrent network state that underlies actual attentive processing in nonhuman primate long-range attention networks (Womelsdorf and Everling 2015).

One important caveat of the burst-network hypothesis that needs to be addressed in future studies is that dendritic burst mechanisms have been described at the cellular level exclusively for pyramidal cells and not interneurons (Larkum et al. 1999). Indeed, for the class of pyramidal cells, fast burst spike events are easier generated in neurons with larger dendritic trees (Mason and Larkman 1990; Yang et al. 1996; van Ooyen and van Elburg 2014). In contrast, we report that not only pyramidal cells, but also narrow spiking neurons that are putative interneurons, fire bursts that relate stronger to network states than their isolated spikes. One intriguing possibility to resolve this conundrum is to assume that some classes of interneurons are endowed with a burst firing mechanisms that co-localizes with the main spike mechanism in the soma and is independent from large dendritic trees (Krahe and Gabbiani 2004). Such co-localized mechanisms exist and are believed to be more likely activated with enhanced barrages of synaptic inputs, characteristic of enhanced network activation (Krahe and Gabbiani 2004). It might thus be possible that bursts characterize enhanced network activity states across neuron classes. It will be important in future studies to characterize the burst firing neuron types more precisely to infer which sub-classes of interneurons participate in attention specific burst firing.

### 3.2 - Putative interneuron bursts, neuronal synchronization and network oscillations

We found that burst firing of putative interneurons is associated with stronger theta power than isolated interneuron spikes. The burst specific power modulation started shortly before the time of the burst event and lasted for ∼100ms after the first burst spike. This finding is significant as it highlights that burst spikes have a unique relation to the strength of theta rhythmic activity of the local network. The acausal nature of this finding - with theta power increasing already shortly before the burst event - indicates not only that bursts are systematically locked to the ongoing low frequency rhythm (Ray and Maunsell 2011), but that they could play a more active role to sustain, facilitate, or even initiate synchronized oscillations in the local neural circuit. Such a role is consistent with recent optogenetic in-vitro studies that implicate interneuron bursts to exert a powerful inhibitory synchronization pulse to the surrounded pyramidal cell network (Berger et al. 2010; Hilscher et al. 2017). In particular, the burst firing of single Martinotti cells have been shown to impose compound inhibitory postsynaptic potentials strong enough to silence connected multiple pyramidal cells in the local circuit (Hilscher et al., 2017, Fig. S6). In these experiments, the burst-induced synchronized inhibition resets pyramidal cell activity, whose action potentials synchronize during the recovery from inhibition (Hilscher et al. 2017). Intriguingly, optogenetically induced rhythmic, low frequency (<20Hz), inhibitory pulsing of these burst firing interneurons not only initiated de-novo synchronized firing, but sustained rhythmically synchronized activation in the nearby pyramidal cell network (Hilscher et al. 2017). These widespread consequences of interneuron bursts have been documented specifically for Martinotti cells located close to layer 5. Modeling studies of the role of inhibitory bursts strongly support the potential of this class of interneuron to initiate and reset ongoing oscillatory activity by systematically silencing asynchronous pyramidal cell firing due to the temporally extended inhibitory potential of the burst spikes (Sahasranamam et al. 2016). It will be an important task for future studies to characterize more precisely which narrow spiking neurons in extracellular recordings may correspond to the Martinotti, low threshold firing cell type. For example, a previous study has suggested that at least three narrow spiking neurons are distinguishable in extracellular recordings with differences in the cells propensity to synchronize at theta versus beta activity (Ardid et al. 2015). In a similar vein, we believe that with sufficient dense recording of layer 5 cell activity it will be possible to test whether bursts of interneuron subclasses are directly initiating or maintaining periodic activity at slow (<20Hz) frequencies in prefrontal brain networks during attention states.

### 3.3 - Beta-synchronized burst firing may facilitate rapid changes in activation states

We found that beta power is more strongly modulated during phase synchronized bursts, rather than nonbursts, for putative interneurons (**Fig. 7B,C**), and that bursts of putative pyramidal cells coincide with strong phase dependent beta power over a shifted phase range of the beta cycle (**Fig. 7E,F**). These findings reveal a novel link of burst specific firing events of ACC/PFC neurons to phase synchronization in the beta frequency band and could provide important constraints for models of beta generation. For example, beta synchronized oscillations in the ACC/PFC are often not sustained, but briefly waxing and waning events of about three consecutive beta cycles (Murthy and Fetz 1996; Feingold et al. 2015; Sherman et al. 2016). These brief beta events have been traced mechanistically to the coincident activation of dendritic and proximal inputs of a network of inhibitory and pyramidal cells (Sherman et al. 2016). One observation of the model is that brief beta events could emerge even if the distal dendritic input is not-rhythmic (Sherman et al. 2016), while the output of the beta generating circuits does carry beta rhythmicity that functionally couples frontal cortex with long-range targets (Cagnan et al. 2015; Feingold et al. 2015). Another observation is that decreasing the variability in the timing of inputs results in greater beta modulation (Sherman et al. 2016). These observations are consistent with two of our findings. First, beta power modulation in the local population surrounding NS cells could be the result of more strongly synchronized, less variable, inputs during bursts, as opposed to nonbursts. Likewise, putative pyramidal cells’ bursts and nonburst events result in similarly high beta power modulation across burst phases spanning a wider phase range in the beta cycle than nonbursts. This result was evident in a leftward shift of the peak of the phase dependent power modulation and could indicate that burst spikes are elicited already at nonoptimal beta phases at a time when overall beta power has reached a sufficient level, as would be expected if it emerges from similarly synchronized inputs arriving at earlier phases (Sherman et al. 2016). Secondly, the burst spike output itself was linked strongly to beta activity in the local circuit as demonstrated here as burst-specific beta phase synchronization - both in putative interneurons and, to a lesser degree, putative pyramidal cells (**Fig. 6B-D**). This is similar to a previous study, demonstrating strong beta synchronization of bursts in one area to the LFP in other distant brain areas in ACC/PFC (Womelsdorf, Ardid, et al. 2014). Taken together, the burst specific phase synchronization effects may reflect facilitated interactions of the local circuit with long-range connected areas by contributing to a more beta synchronized spike output of the local circuit. This scenario predicts that long-range coherent network activity is supported by mechanisms generating burst firing of single neurons during attention states (Larkum 2013).

### 3.4 - Burst spikes may actively contribute to the local field

Beyond a role of burst firing for network activity inferred from power modulation and phase synchrony, burst firing mechanisms may also directly contribute to the local electrical field activity measured from sharp extracellular electrodes (Sanchez-Vives and McCormick 2000; Buzsáki et al. 2012a; Einevoll et al. 2013). We believe that such a potential direct influence on the LFP should not be understood as a confound, but as an important, possible window into the cellular mechanisms of local circuit formation. In particular, burst firing has been documented in-vivo to affect the extracellular local field by causing post-burst after-hyperpolarization of membranes of the burst firing neurons (discussed in (Buzsáki et al. 2012b)). In pyramidal cells such burst induced hyperpolarization of cell membranes is mediated by Ca++ currents that can increase repolarizing K+ conductances in the somatic region (Hotson and Prince 1980), or by activation of NMDA or Na+ spikes within dendritic compartments, leaving >15ms long traces of hyperpolarization (Nevian et al. 2007; Sjöström et al. 2008). It has been established that NMDA receptors in dendritic spines sense glutamate excitation and have a decay time constant in an estimated range of 10-100ms (Major et al. 2013). This NMDA decay time constant may relate to the finding in (rodent) mPFC slices that localized dendritic glutamate release not only triggers somatic burst firing, but also ≥ 100ms plateau potentials (Milojkovic et al. 2004).

Importantly, these burst related local dendritic effects of longer lasting after-hyperpolarization may affect the local extracellular field when bursts are coordinated in time (Buzsaki et al. 1988; Sanchez-Vives and McCormick 2000; Buzsáki et al. 2012a). This suggestion is consistent with our result showing that bursts spike are significantly coordinated with population-level synchronized theta and beta band activities. We thus speculate that one component of the burst triggered LFP average may be attributable to a longer lasting (∼100ms) hyperpolarization across burst firing neurons in the neural network. It will be an important question for future work to separate such direct field effects from more indirect interactions with the field activity, similar to what has been started in visual cortex (Mitzdorf 1985; Nauhaus et al. 2009; Ray and Maunsell 2011; Teleńczuk et al. 2016).

### 3.5 - Outlook: Functional implications of burst specific network activity

Recent models suggest that burst firing neurons connect to each other during synchronous burst firing to enable efficient synaptic strengthening among those neurons receiving similar dendritic inputs at similar times (Sjöström et al. 2008; Kaifosh and Losonczy 2016; Wilmes et al. 2016). Such coincident activation of neurons switches on long-term potentiation mechanisms and could thus induce plasticity among synapses of those neurons participating in synchronous burst firing. According to this scenario, burst events might have a special role in the formation of networks of neurons participating in the same functional process. Our findings add to this suggestion by showing that burst events can be more strongly modulated by population level rhythmic activities than nonburst singleton events. This enhanced burst-LFP relationship was not only evident specifically for bursts during the attentional state in prefrontal and anterior cingulate cortex, but became evident at those frequency bands that have been most prominently related to endogenously controlled, goal directed behaviors (Larkum 2013; Fries 2015; Womelsdorf and Everling 2015). Taken together, we believe that mechanisms underlying burst spike generation will provide a direct window into the origin of cellular control of higher order attentional behaviors.

## Acknowledgments

This research was supported by grants from the Canadian Institutes of Health Research (CIHR), the Natural Sciences and Engineering Research Council of Canada (NSERC) and the Ontario Ministry of Economic Development and Innovation (MEDI). We thank Stefan Everling for help with the electrophysiological recording and the surgical procedures, Dr. Daniel Kaping for invaluable help with the electrophysiological recording and Johanna Stucke for exceptional animal care. The funders had no role in study design, data collection and analysis, decision to publish, or preparation of the manuscript.

